# Thiosemicarbazone T2 suppresses WNT/β-catenin signaling and limits progression to invasive disease in a mouse intraductal model of triple-negative breast cancer

**DOI:** 10.64898/2026.06.05.730419

**Authors:** Aldana M Sólimo, Marianela Sciacca, Florencia Cascardo, Liliana Finkielsztein, Ana María Eiján, Catalina Lodillinsky, Mariana A Callero

## Abstract

T2, an N4-aryl-substituted thiosemicarbazone, has previously been shown to exert cytotoxic and anti-invasive effects in triple-negative breast cancer (TNBC) and to increase expression of the metastasis suppressor N-myc downstream-regulated gene 1 (NDRG1). Given the role of NDRG1 in regulating epithelial–mesenchymal transition (EMT) and WNT/β-catenin signaling, we investigated the contribution of this pathway to the anti-invasive activity of T2.

The effects of T2 on WNT/β-catenin signaling and associated microRNAs (miR-182-5p and miR-200c) were evaluated in 4T1 cells. In vivo activity was assessed using a fully immunocompetent intraductal 4T1 mouse model that recapitulates the progression from ductal carcinoma in situ (DCIS) to invasive ductal carcinoma (IDC). Tumor progression, invasion, NDRG1 expression, and WNT/β-catenin pathway components were analyzed.

T2 reduced WNT/β-catenin signaling and modulated the expression of miR-182-5p and miR-200c in vitro. In the MIND model, T2 decreased the frequency of invasive lesions and reduced β-catenin, ZEB1, and c-Myc expression while increasing NDRG1 levels. β-catenin localization differed between lesion types, showing predominantly membrane-associated staining in DCIS lesions and a diffuse cytoplasmic distribution in invasive foci.

These findings identify WNT/β-catenin signaling and NDRG1-associated pathways as potential mediators of the anti-invasive effects of T2 in TNBC. The reduction in invasive progression observed in the MIND model supports further investigation of this compound in preclinical models of TNBC.

## INTRODUCTION

Breast cancer (BC) remains the most frequently diagnosed malignancy and the leading cause of cancer-related death among women worldwide [1–3]. Based on the expression of estrogen receptor (ER), progesterone receptor (PR), and human epidermal growth factor receptor 2 (HER2), BC comprises biologically distinct subtypes with different clinical behaviors and therapeutic vulnerabilities. Among them, triple-negative breast cancer (TNBC), defined by the absence of ER, PR, and HER2 expression, is associated with aggressive clinical behavior, early recurrence, and poor prognosis [4,5]. Owing to the lack of actionable molecular targets, chemotherapy remains the main systemic treatment option for most patients with TNBC.

Thiosemicarbazones (TSCs) are synthetic compounds with metal-chelating properties that have attracted attention as anticancer agents. By binding transition metals, particularly iron, TSCs can disrupt cellular iron homeostasis, promote oxidative stress, and interfere with signaling pathways involved in tumor progression [6].

Preclinical evaluation of novel therapeutic strategies requires experimental models that reproduce key features of human disease. Although orthotopic and subcutaneous transplantation models are widely used in breast cancer research [7], they do not fully recapitulate the natural progression of ductal tumors. In patients, invasive ductal carcinoma (IDC) typically arises from pre-existing ductal carcinoma in situ (DCIS) lesions through progressive invasion of the surrounding mammary tissue [8]. The mouse intraductal (MIND) model reproduces this sequence of events by allowing tumor cells to grow within the mammary ducts before invading the adjacent stroma, thereby providing a physiologically relevant system to study early breast cancer progression [9].

In a previous study, we demonstrated that T2, an N4-aryl-substituted thiosemicarbazone, exerts cytotoxic, anti-invasive, and anti-metastatic effects in 4T1 TNBC cells both in vitro and in vivo [10]. These effects were accompanied by increased E-cadherin expression, reduced α-SMA levels, cortical actin organization, and induction of a more differentiated epithelial phenotype. We also observed increased NDRG1 protein expression without changes in NDRG1 mRNA levels NDRG1 is a well-established metastasis suppressor that negatively regulates several pathways implicated in tumor progression, including NF-κB, EGFR, E-cadherin, and WNT/β-catenin signaling [11]. Given the central role of WNT/β-catenin signaling in TNBC progression and epithelial–mesenchymal transition (EMT), we investigated whether this pathway contributes to the antitumor activity of T2. We further examined the expression of microRNAs associated with NDRG1 and WNT signaling and evaluated the effects of T2 in a fully immunocompetent 4T1 MIND model.

Our results show that T2 suppresses WNT/β-catenin signaling, modulates miR-182-5p and miR-200c expression, and reduces the progression of in situ lesions toward invasive disease. In the MIND model, these effects were accompanied by decreased β-catenin, ZEB1, and c-Myc expression together with increased NDRG1 levels. Collectively, these findings support a role for NDRG1-associated regulation of WNT/β-catenin signaling in the antitumor activity of T2.

## MATERIALS AND METHODS

### Cell culture

The 4T1 mouse mammary carcinoma cell line was obtained from ATCC and cultured in RPMI medium supplemented with 10% FBS and 80 μg/ml gentamicin at 37°C in 5% CO_2_. Cells were tested for mycoplasma contamination by PCR at least once a month.

### qRT-PCR

Cells were treated with 0.2% DMSO or 5 µM T2 for 24 h, with 10 mM LiCl added to induce Wnt signaling (10 mM NaCl as control). RNA was extracted using BIO-ZOL (PBL, Argentina). Two micrograms of RNA were reverse transcribed with M-MLV and random primers. qPCR for Axin2, CD44, c-Myc, TCF-7, Snail, Twist, and Vimentin was performed using TransStar Green on a Biorad C1000 Thermal Cycler (primer sequences in Supplementary Table 1). Cycling included 40 cycles at 94°C, 60°C, and 72°C (30 s each) with fluorescence measured after elongation. Expression was normalized to HPRT and analyzed by the 2^-ΔΔCt method. Samples were run in duplicate with no-template controls.

### Stem-Loop PCR[12]

Briefly, cells were treated with 0.2% DMSO or T2 5 µM for 24 h. RNA extraction and reverse transcription were performed as before, using 1 µM stem-loop primers (sequences in Supplementary Table 2) instead of random primers. qPCR used specific forward primers (Supplementary Table 2) and a universal reverse primer. Expression was normalized to U6 snRNA and analyzed by the 2^-ΔΔCt method. Samples were run in duplicate with no-template controls.

### Western Blot

Proteins were extracted with RIPA-EDTA buffer from confluent 4T1 cell monolayers treated with 0.2% DMSO, 5 µM T2 plus 10 mM LiCl, or NaCl for 24 h. Protein concentration was measured by Bradford assay.

Proteins were separated by SDS-PAGE and transferred to PVDF membranes. Membranes were blocked and then incubated with primary antibodies: GAPDH (1:1000, Novus NB300-327), β-catenin (1:1000, Cell Signaling #8480), ZEB1/TCF8 (1:250, Cell Signaling #3396), and c-Myc (1:500, Cell Signaling #D84C12). HRP-conjugated secondary antibodies (mouse and rabbit, 1:500, Novus HAF007/HAF008) were applied.

### Transfection

Plasmids M50 Super 8x TOPFlash (Addgene, #12456) [13] and Renilla-luciferase (pRL-CMV, Promega) were transfected using FUGENE HD (Promega). After 24h, cells were treated with DMSO or T2 5 µM in combination with 3 ng/ml TGF-β for another 24h. Luminescence was measured using the Dual-Luciferase Reporter Assay System kit (Promega).

### Animals

BALB/c mice (12-16 weeks old) were obtained and housed in the animal facility at the Angel H. Roffo Oncology Institute. Experimental protocols were reviewed and approved by the institute’s animal use committee (protocol 2017/01).

### Intraductal Inoculation and Whole Mount

Eleven to thirteen-week-old virgin female mice were anesthetized (xylazine and ketamine) and abdominally shaved. Nipples of both fourth mammary glands were snipped, and 2 µl of cell suspension (17,500 or 7,500 cells/µl) was injected into each gland using a 30-gauge Hamilton syringe with a blunt ½-inch needle. Treated mice (6 per group) received T2 (25 mg/kg) or vehicle (30% propylene glycol, 17% DMSO in saline) via five intraperitoneal doses starting two days post-inoculation, every 48 h. Mice were euthanized 7 or 14 days later, mammary glands harvested, spread on slides, and fixed overnight in methanol, chloroform, and acetic acid (methacarn). Slides were washed with 70% ethanol and water, stained with 0.2% carmine and 0.5% potassium aluminum sulfate for 1–2 days, dehydrated through ethanol and xylene, and photographed under a stereoscope before paraffin embedding for sectioning and immunofluorescence or hematoxylin-eosin (HE) staining.

### Immunofluorescence

Slides were deparaffinized and rehydrated. For antigen retrieval, slides were incubated in a citric acid/sodium citrate buffer heated at 100°C for 8 min. For permeabilization, slides were incubated twice with 0.1% Triton, 1% FBS in PBS for 10 min, followed by blocking with 5% FBS for 1 h. Slides were incubated overnight with the primary antibodies at 4°C, then washed and incubated with corresponding secondary antibodies for 2 h at room temperature. DAPI stain was applied for 15 minutes, and slides were mounted with Mowiol (Calbiochem). Images were obtained using a Nikon ECLIPSE E400 microscope and processed with ImageJ software.

### Statistical analysis

In vitro assays were carried out in three independent experiments (n = 3). The in vivo experiment was conducted once in accordance with our institute’s animal use committee’s reduction principle, using six animals per group, with both mammary glands from each animal evaluated separately. Data were analyzed using GraphPad Prism 5.0. Two-group comparisons employed Student’s t-test with Welch’s correction for unequal variances. For comparisons of more than two groups, the Kruskal-Wallis test with Dunn’s post hoc test was used. Statistical significance was set at p < 0.05.

### Data Avaibility

The datasets generated during and/or analyzed during the current study are available from the corresponding author on reasonable request.

### AI use

The authors used ChatGPT (OpenAI) to assist in improving the clarity and readability of the manuscript. All content, data interpretation, and conclusions were conceived, verified, and approved by the authors.

## RESULTS

### T2 downmodulates EMT markers and WNT/β-catenin signaling

Since T2 promoted a more differentiated epithelial phenotype [14], we investigated whether it could reverse epithelial-mesenchymal transition (EMT) in 4T1 cells. T2 treatment (5 µM) reduced the expression of Snail, Twist, and Vimentin in LiCl-treated cells (Figure 1A) [15]. A similar effect was observed for the WNT/β-catenin target genes Axin2, CD44, c-Myc, and TCF-7 (Figure 1B). Western blot confirmed significantly decreased protein levels of the Wnt/β-catenin effector ZEB1, while c-Myc, and β-catenin showed non-significant modulation (Figures 1C–D). TOPFlash reporter assays showed reduced β-catenin transcriptional activity in T2-treated cells following TGF-β stimulation (Figure 1E).

**Fig. 1.**
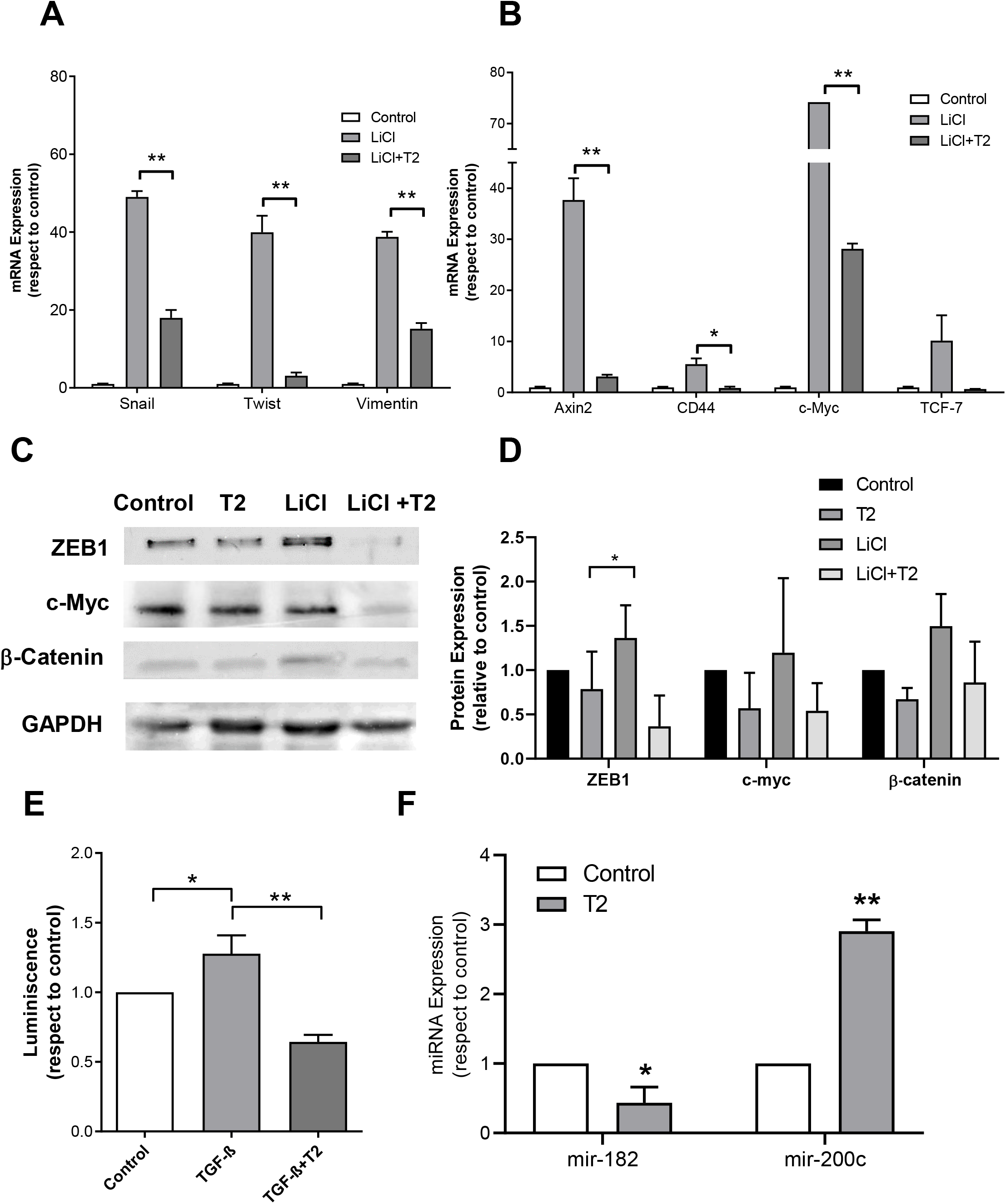
Evaluation of EMT markers and Wnt/β-catenin signaling pathway in 4T1 cells. LiCl-induced EMT cells were treated with DMSO (control) or T2 5 µM for 24 h. (A) Expression of EMT markers and (B) Wnt/β-catenin molecular targets analyzed by Real Time PCR. (C) Western blots illustrating ZEB1, c-Myc, and β-catenin protein expression. A representative blot of three is shown. (D) Densitometric analyses of three independent western blots. (E) Luciferase activity in cells previously transfected with TOPFlash TCF/LEF and Renilla reporter and subsequently treated with DMSO (control), TGF-β (Wnt/β-catenin inducer), or TGF-β +T2 (5 µM) for 24 hours. (F) Cells were treated with DMSO (control) or with T2 5 μM. mir-182-5p and mir-200c expression evaluated by Stem-loop PCR. Results are expressed as mean ± SD. *p < 0.05, **p < 0.01 (n = 3)

### T2-mediated regulation of NDRG1 and Wnt/β-catenin signaling is associated with microRNA modulation

Previously, we showed that T2 affected NDRG1 protein but not mRNA levels. To further explore its role in EMT reversal, we analyzed miR-182-5p, which downregulates NDRG1 and enhances Wnt/β-catenin signaling [16,17], and miR-200c [18], known to inhibit this pathway. Stem-loop qPCR revealed that T2 downregulated miR-182-5p and upregulated miR-200c (Figure 1F), suggesting that both are molecular targets of T2.

### Optimization of 4T1 TNBC MIND Model

To evaluate the effects of T2 during early tumor progression, a 4T1 MIND model was established. This model reproduces the transition from DCIS to IDC observed in human breast cancer and provides a suitable platform to investigate mechanisms associated with local invasion [19–22]. Initially, 35,000 4T1 cells were injected intraductally, resulting in rapid and extensive tumor growth (Supp Figure 1A). Reducing the inoculum to 15,000 cells yielded smaller foci at 7 days and distinct DCIS and IDC stages at 14 days, while preserving ductal architecture. Therefore, we decided to continue our experiments with 15.000 cells. Upon histological analysis we classified ducts as empty, DCIS, or IDC. A significant increase in IDC foci and a concurrent decrease in DCIS were observed at 14 days compared to 7 days, with no change in empty ducts (Supp Figure 1B-C). Immunofluorescence showed vimentin expression in both stages without significant differences (Figure 2A-B). Proliferation (PCNA/DAPI) increased significantly at 14 days (29±7%) versus 7 days (12±4%) (Figure 2C).

**Fig. 2.**
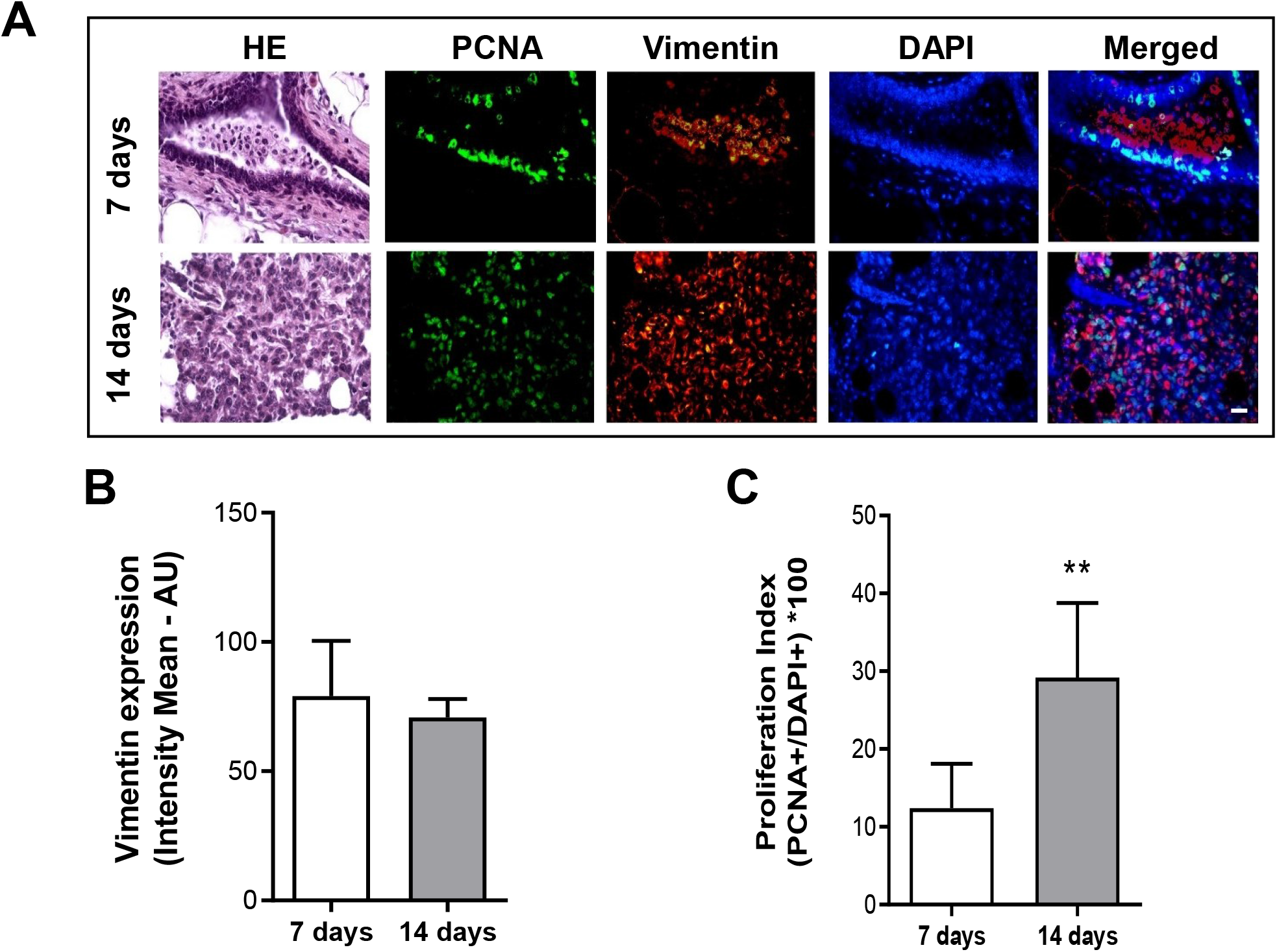
Characterization of 4T1 MIND Model. Immunofluorescence of BALB/c mouse mammary gland removed after 7 or 14 days following intraductal 4T1 cell inoculation. (A) Representative images of proliferation marker PCNA (green), mesenchymal marker vimentin (red), and DAPI staining (blue); magnification 400x. Magnification bar: 50µm. (B) Vimentin expression calculated as the percentage of positive red staining with respect to positive blue staining. (C) Proliferation index calculated as the percentage of positive green staining with respect to positive blue staining. Data represent mean ± SD from an experiment with 6 mice per group; ** *p*<0.01 vs. 7 days (Student’s t-test)

Pan-cytokeratin and α-SMA staining confirmed epithelial tumor identity and loss of ductal integrity at invasive lesions (Supp Figure 2A). Immune profiling revealed a significant drop in the CD4+/CD8+ ratio (from 4.5±2 to 1.4±0.6; p<0.05) due to increased CD8+ cells, consistent with tumor progression (Supplementary Figure 2B).

### T2 downmodulates in situ-invasive transition in the 4T1 MIND model

BALB/c mice were intraductally injected with 4T1 cells, and 48 hours later, received T2 treatment (5 doses of 25 mg/kg IP every other day) as in previous studies [14]. Mice were sacrificed 14 days post-inoculation for analysis.

The whole mount examination showed no major morphological differences between treated and control glands (Supplementary Figure 3). However, histology revealed a significant reduction in IDC ducts in T2-treated mice (13±1% vs. 20±3%, p<0.05; Figures 3A-B). Immunofluorescence for PCNA indicated decreased tumor cell proliferation with T2 (27±6% vs. 43±4%, p<0.05; Figure 3C), while vimentin expression remained unchanged (Figure 4D).

**Fig. 3.**
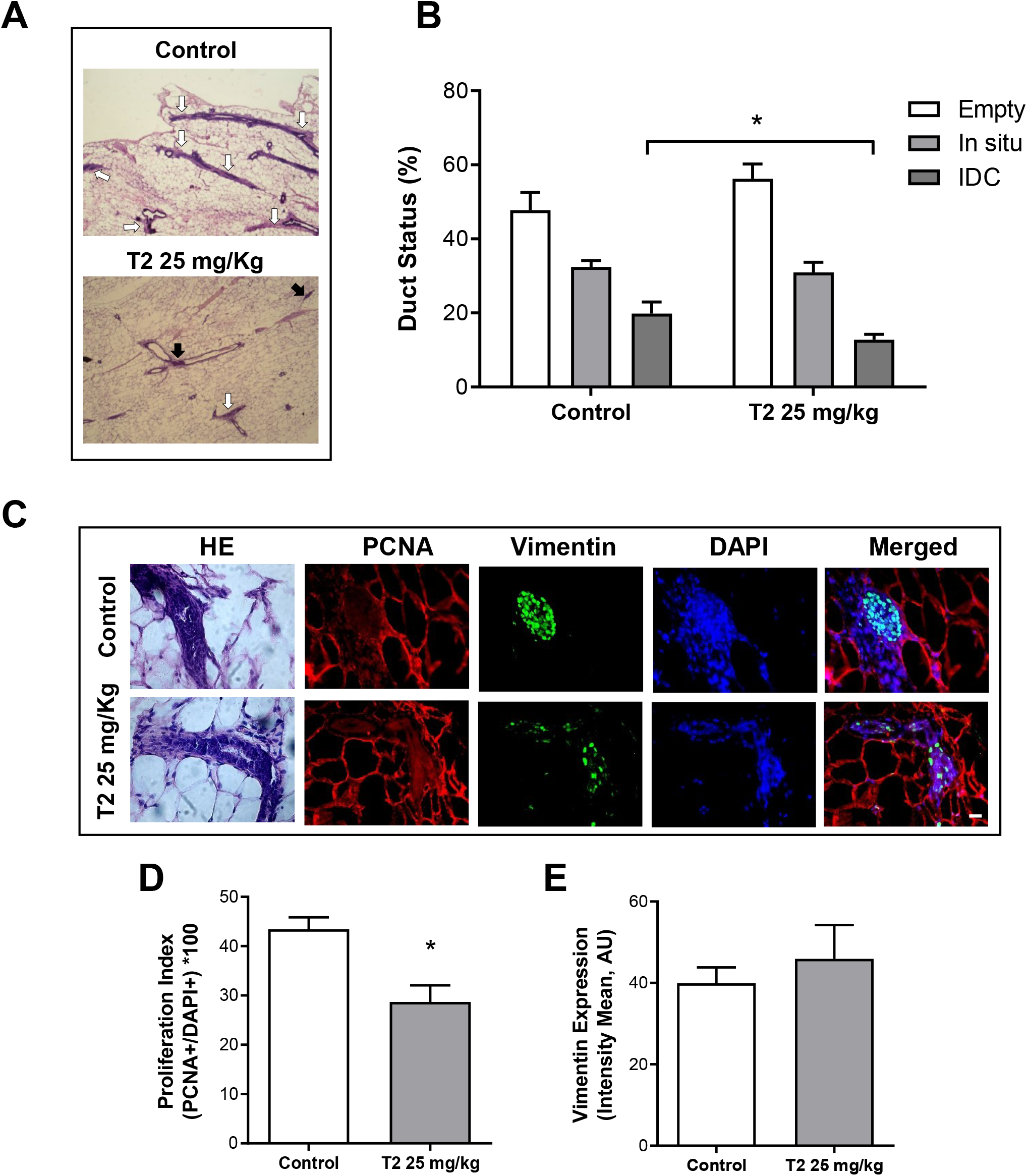
T2 action on the 4T1 invasive capacity in the MIND model. (A) BALB/c mouse breast extracted 14 days after inoculating 15,000 4T1 cells and administering five intraperitoneal doses of either vehicle (control) or T2 at 25 mg/kg. (A) Representative micrographs of the hematoxylin-eosin staining of the mammary gland; DCIS: black arrows, IDC: white arrows (400x magnification). (B) Duct status quantification (empty, with DCIS, or IDC growth) observed in sections stained with hematoxylin-eosin. The mean and standard error of an experiment with 6 mice per group are shown, *p<0.05 respect to control. (C) Representative images of proliferation marker PCNA (green), mesenchymal marker vimentin (red), and DAPI staining (blue); magnification 400x. Magnification bar: 50μm. (D) Proliferation Index calculated as the percentage of positive green staining with respect to positive blue staining. (E) Vimentin expression calculated as the percentage of positive red staining with respect to positive blue staining (5-10 sections per animal, n=3 per group).

**Fig. 4.**
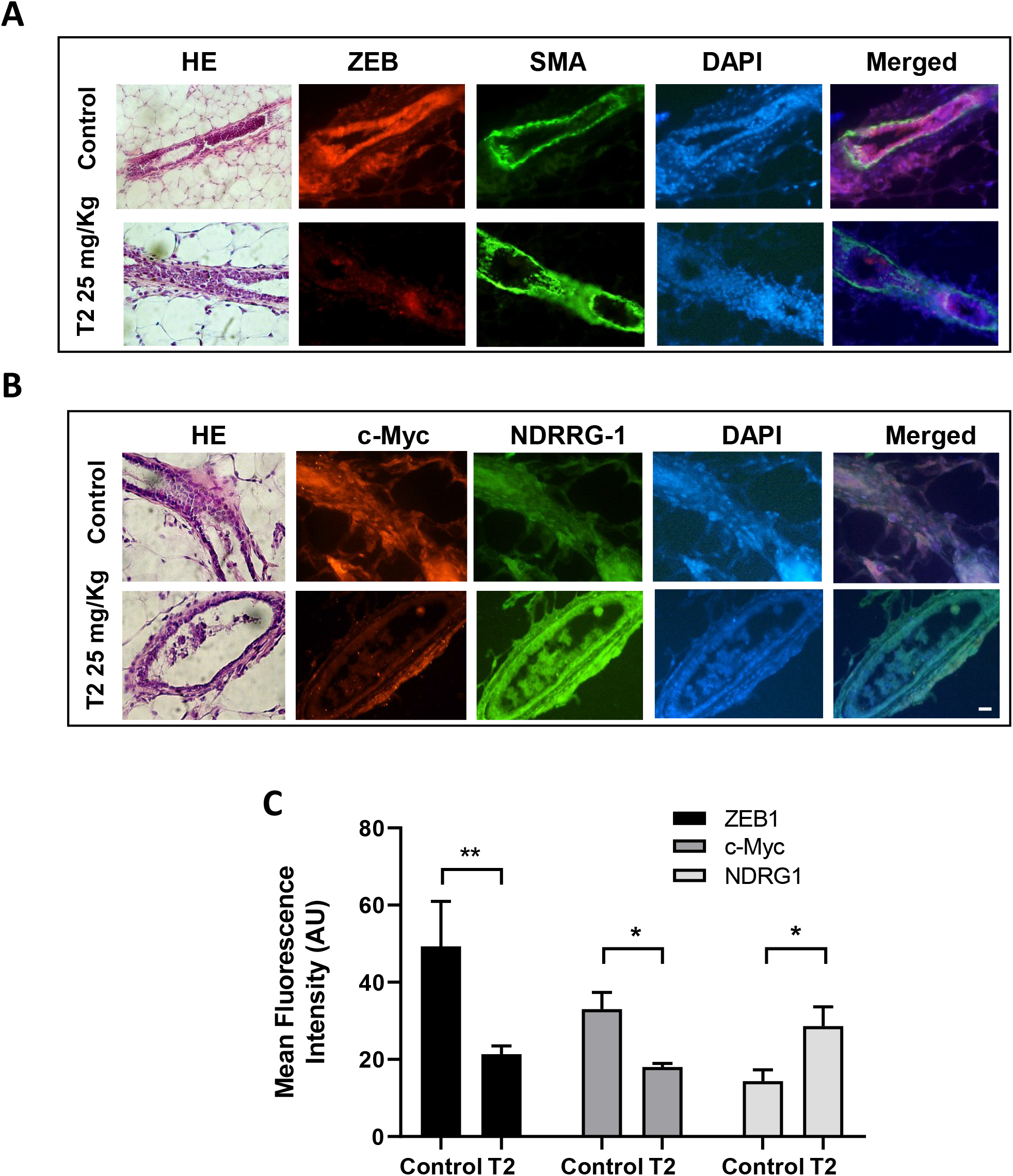
T2 effects on ZEB1, c-Myc, and NDRG1 expression in the 4T1 MIND model. (A) Representative immunofluorescence images for α-SMA (green) and ZEB1 (red) with DAPI (blue) performed on mammary gland sections from mice inoculated with 4T1 cells and treated with 25 mg/kg T2 or vehicle. (B) Representative HE stained and immunofluorescence images for NDRG1 (green) and c-Myc (red) with DAPI (blue) performed on mammary gland sections from mice inoculated with 4T1 cells and treated with 25 mg/kg T2 or vehicle. Magnification 400x. Magnification bar: 50μm. (C) Quantification of ZEB1, c-Myc, and NDRG1 expression evaluated as red or green mean fluorescence intensity on DCIS or IDC foci (5-10 sections per animal, n=3 per group). Data represent mean ± SD from an experiment with 6 animals per group; *p*<0.05 vs. control (Student’s t-test)

### T2 reduces ZEB1, c-Myc, and β-catenin expression while increasing NDRG1 level in the 4T1 MIND model

Based on the in vitro modulation of NDRG1 and WNT/β-catenin signaling, immunofluorescence analysis was performed to evaluate ZEB1, c-Myc, NDRG1, and β-catenin expression in mammary gland sections. T2 treatment decreased ZEB1 and c-Myc levels (AU: 49±7 vs. 21±1 and 33±3 vs. 18±1, respectively; p<0.05) and increased NDRG1 fluorescence (AU: 14±2 vs. 29±3; p<0.05) in 4T1 tumor foci, irrespective of lesion type (Figure 4A-B). β-catenin localization differed between DCIS and IDC lesions. In control tumors, β-catenin staining was predominantly membrane-associated in DCIS lesions, whereas invasive foci showed reduced membrane localization and a diffuse cytoplasmic distribution (Figure 5A). T2 treatment reduced β-catenin expression in both compartments, from 60±1 to 36±6 AU in DCIS lesions (p<0.01) and from 44±8 to 20±4 AU in IDC lesions (p<0.05) (Figure 5B).

**Fig. 5.**
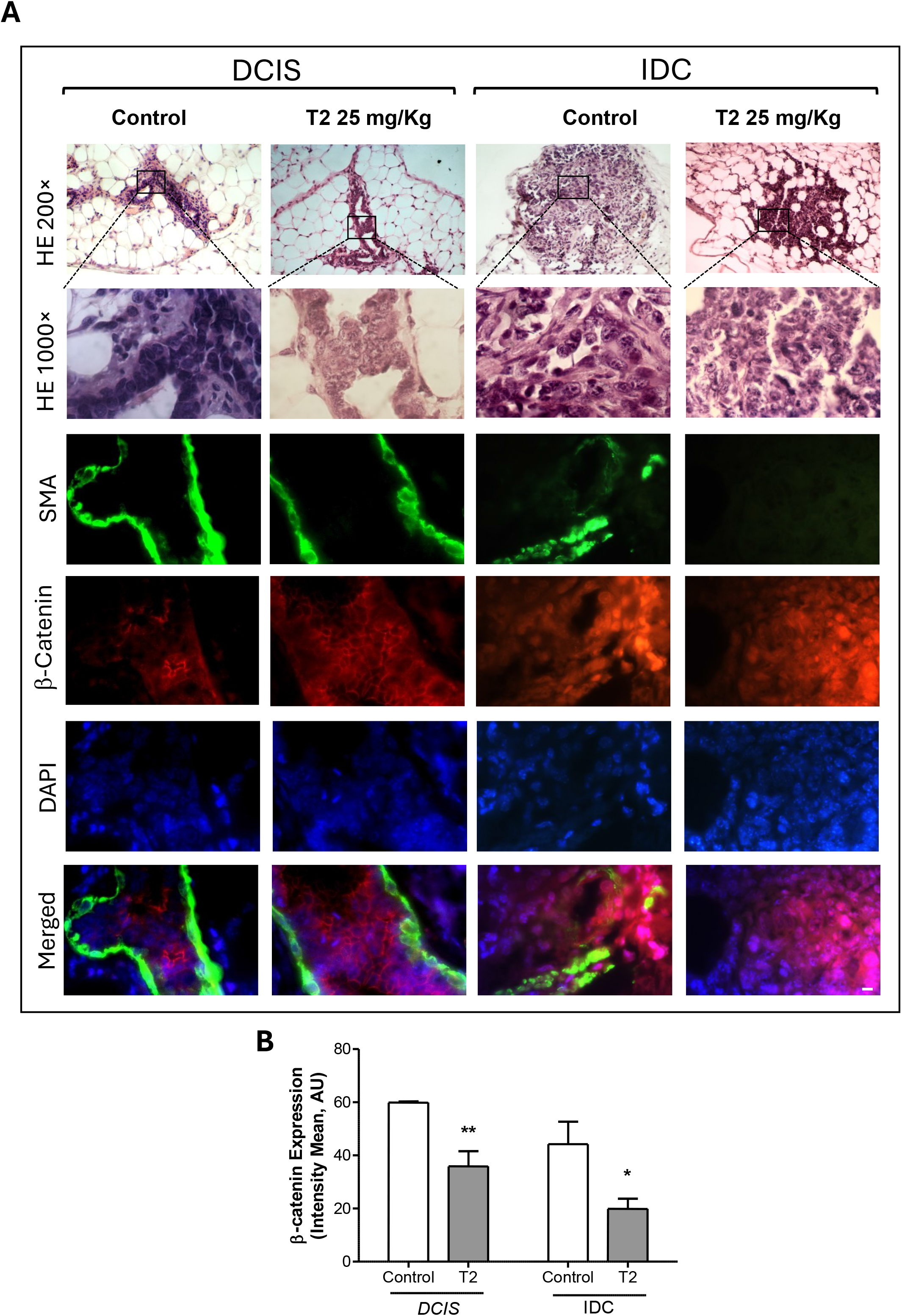
T2 effects on β-catenin expression and localization in the 4T1 MIND model. Representative HE-stained and immunofluorescence images for α-SMA (green) and β-catenin (red) with DAPI (blue) performed on mammary gland sections from mice inoculated with 4T1 cells and treated with 25 mg/kg T2 or vehicle. For each condition, a field of 4T1 cells in situ and a field of 4T1 cells that invaded the glandular stroma are shown, 200× and 1000×. Magnification bar: 10μm. B: Quantification of A, Mean and standard error of an experiment with 6 animals per group are shown, *p<0.05 **p<0.01. (5-10 sections per animal, n=3 per group).

## DISCUSSION

In this study, we show that the thiosemicarbazone derivative T2 modulates WNT/β-catenin signaling, regulates EMT-associated factors, and reduces invasive progression in a 4T1 MIND model of triple-negative breast cancer. T2 decreased the expression of β-catenin target genes, reduced β-catenin transcriptional activity, altered the expression of EMT regulators, and reduced the frequency of invasive lesions in vivo.

Aberrant activation of WNT/β-catenin signaling contributes to TNBC progression by promoting proliferation, stemness, invasion, and metastatic dissemination [23–27] Consistent with this role, T2 reduced the expression of several WNT/β-catenin downstream targets, including c-Myc, CD44, Axin2, and TCF7, and decreased β-catenin-dependent transcriptional activity. T2 also reduced the expression of EMT-associated transcription factors such as Snail, Twist, and ZEB1, supporting an effect on cellular programs linked to tumor plasticity and invasion [28,29].

Interestingly, total β-catenin protein levels were only modestly affected in vitro, whereas TOPFlash assays revealed a significant reduction in β-catenin transcriptional activity. These findings suggest that T2 may preferentially affect the transcriptionally active fraction of β-catenin rather than its overall abundance. In agreement with this possibility, NDRG1 has been reported to suppress β-catenin signaling by limiting its nuclear activity through mechanisms that do not necessarily require a reduction in total β-catenin protein levels [30]

NDRG1 emerged as a central component of the response to T2. In our previous work, T2 increased NDRG1 protein expression without affecting NDRG1 mRNA levels [10]. Here, this observation was accompanied by reduced miR-182-5p expression and increased miR-200c levels. miR-182-5p has been associated with enhanced β-catenin signaling and repression of NDRG1 expression [31–33], whereas miR-200c is a well-established inhibitor of EMT through the regulation of ZEB family transcription factors[34–36]. Although the direct contribution of these microRNAs to the effects of T2 remains to be determined, their modulation is consistent with the observed changes in NDRG1, ZEB1, and WNT/β-catenin signaling.

The lack of significant changes in vimentin expression after T2 treatment, despite the modulation of other EMT-related markers, may reflect the dynamic and heterogeneous nature of the EMT process. Rather than a binary transition between epithelial and mesenchymal states, EMT is currently viewed as a continuum in which cells acquire intermediate phenotypes characterized by distinct combinations of epithelial and mesenchymal traits [37]. Within this framework, transcriptional regulators such as Snail and ZEB1 are frequently altered before changes become evident in structural markers such as vimentin. The simultaneous reduction of ZEB1, c-Myc, and β-catenin in the absence of detectable changes in vimentin expression is therefore compatible with a partial shift away from a mesenchymal phenotype rather than a complete phenotypic reversion [37]

The in vivo findings support the biological relevance of these molecular changes. Using the MIND model, which reproduces the progression from DCIS to IDC in an immunocompetent setting [38,39], we observed a reduction in invasive lesions and decreased tumor cell proliferation following T2 treatment. In addition, β-catenin localization differed between lesion types. DCIS lesions displayed predominantly membrane-associated β-catenin, whereas invasive foci showed loss of membrane localization and a diffuse cytoplasmic distribution. This pattern is consistent with the disruption of epithelial cell-cell adhesion that accompanies local invasion and progression toward IDC [39]T2 reduced β-catenin expression in both DCIS and IDC lesions while increasing NDRG1 expression and decreasing ZEB1 and c-Myc levels.

Previous transcriptomic and pathological studies have identified EMT-associated programs as important drivers of progression from DCIS to invasive disease [40]Likewise, elevated expression of EMT-related factors, including ZEB1 and other mesenchymal regulators, has been associated with poor clinical outcome in TNBC [27,28]. Our findings place T2 within this biological context and support the possibility that inhibition of WNT/β-catenin-dependent programs contributes to its anti-invasive activity.

Like other thiosemicarbazones, T2 promotes oxidative stress and reactive oxygen species generation [10]. These effects may contribute to the modulation of WNT/β-catenin signaling, either directly through redox-sensitive pathways or indirectly through induction of NDRG1 expression [29,30,41]. The relative contribution of these mechanisms remains unclear and warrants further investigation. Future studies should also address the pharmacological properties of T2 and evaluate its activity in additional preclinical models, including patient-derived systems.

Overall, our data identify WNT/β-catenin signaling and NDRG1-associated pathways as potential mediators of the anti-invasive effects of T2 in TNBC.

## Conclusion

T2 modulated WNT/β-catenin signaling, increased NDRG1 expression, altered the expression of EMT-associated microRNAs, and reduced the progression of DCIS lesions toward invasive disease in the 4T1 MIND model. These findings identify WNT/β-catenin and NDRG1-associated pathways as potential mediators of the anti-invasive activity of T2 and support further investigation of this compound in preclinical models of TNBC.

---

**Suppl. Table 1.**
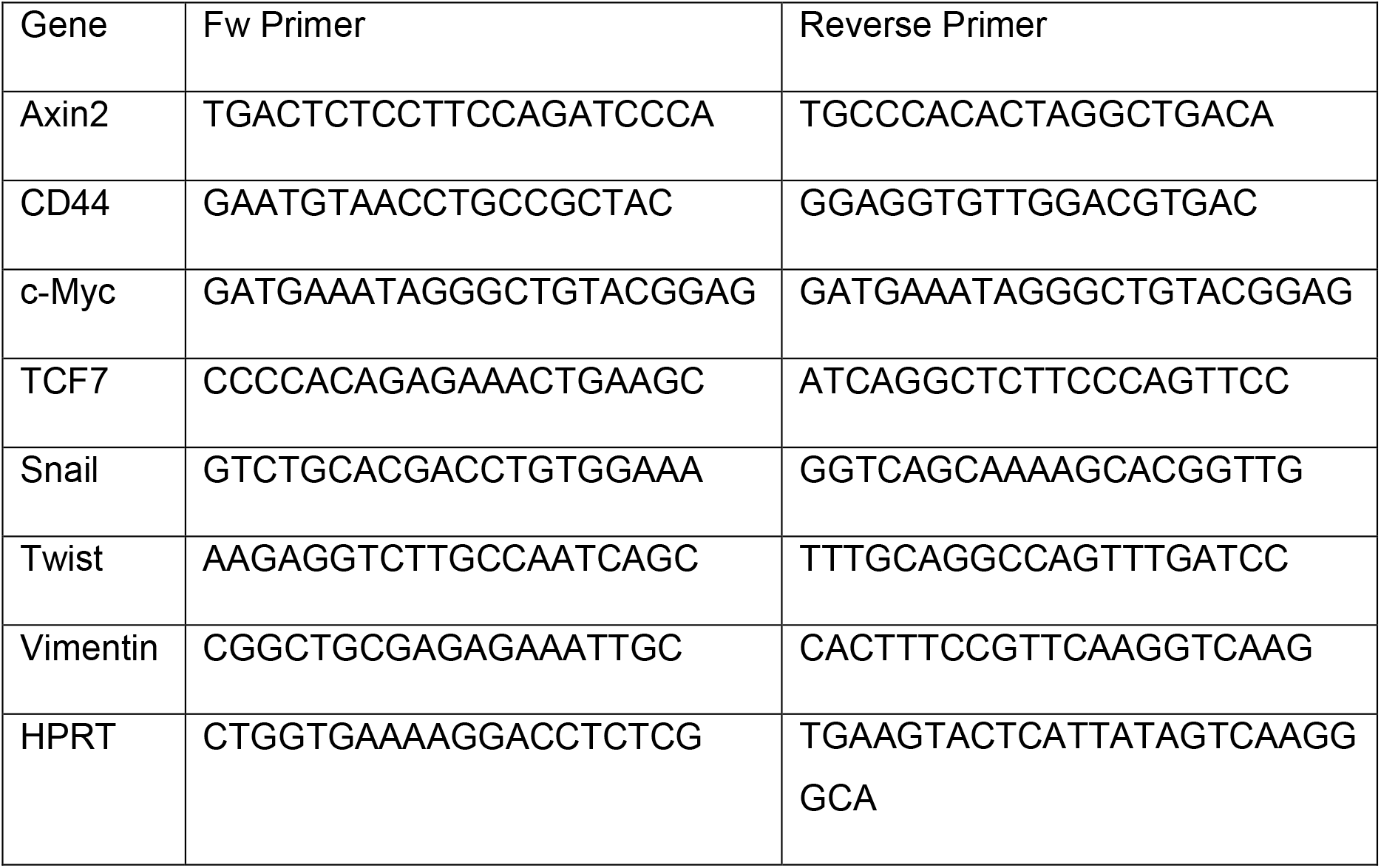
qPCR primer sequences for mRNA quantification.

**Suppl. Table 2.**
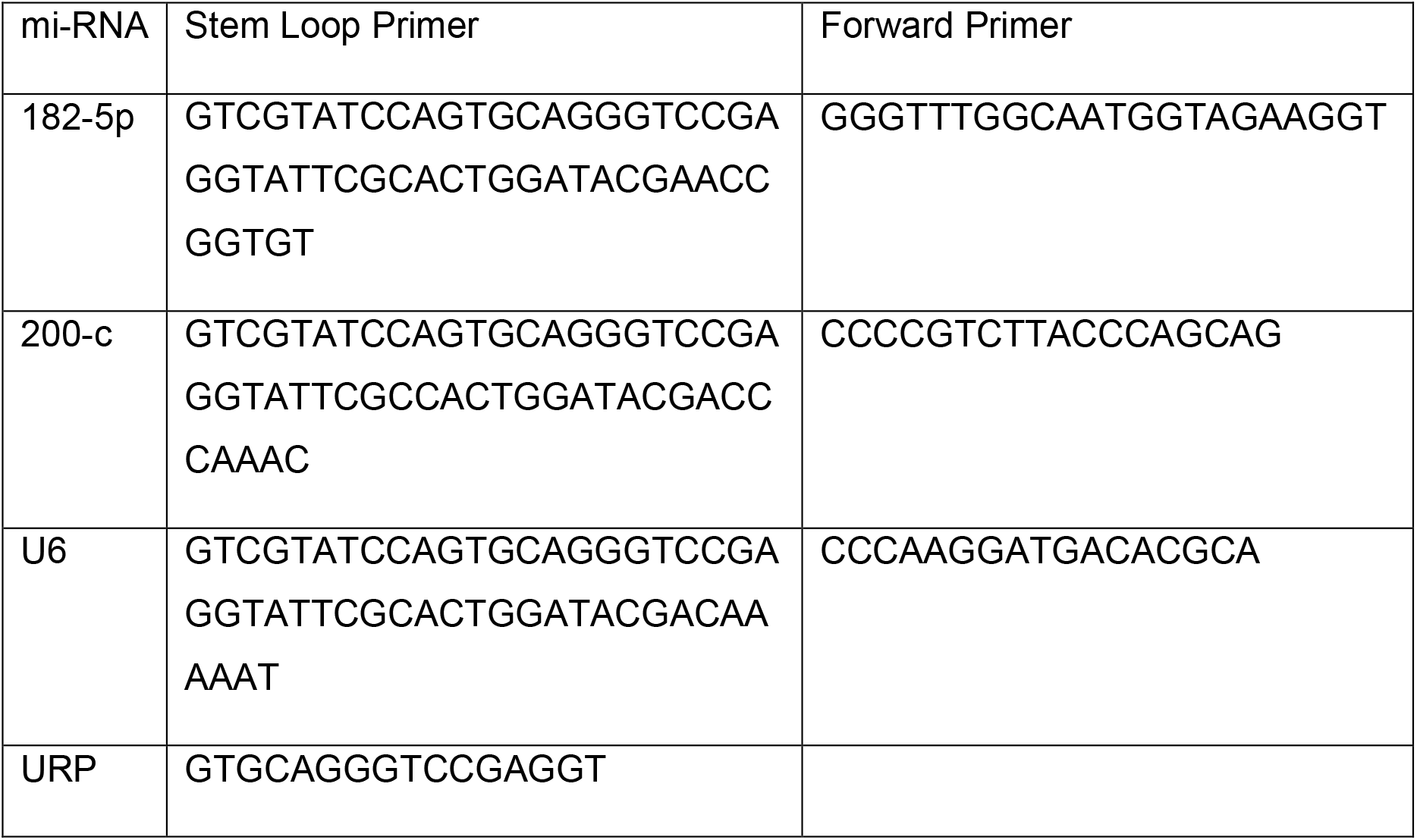
Stem-loop and forward primer sequences for mi-RNA quantification. URP: Universal Reverse Primer.

