## Supplementary figures and images for "Thiosemicarbazone T2 suppresses WNT/β-catenin signaling and limits progression to invasive disease in a mouse intraductal model of triple-negative breast cancer"

### Supplemental Figures

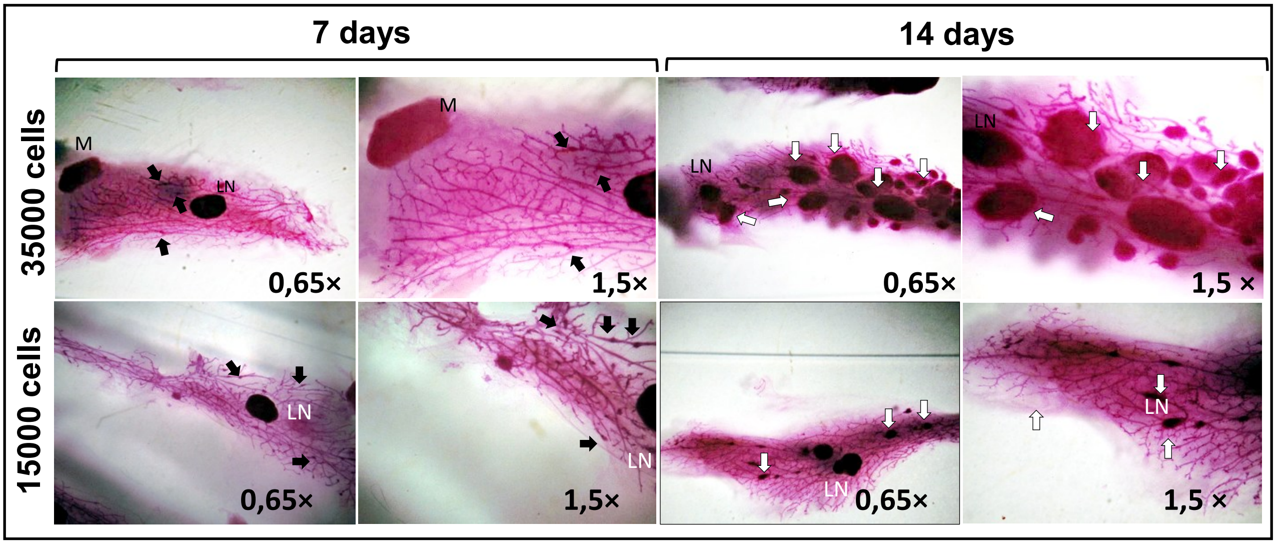

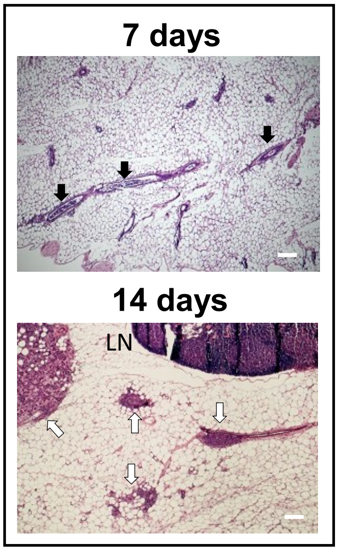

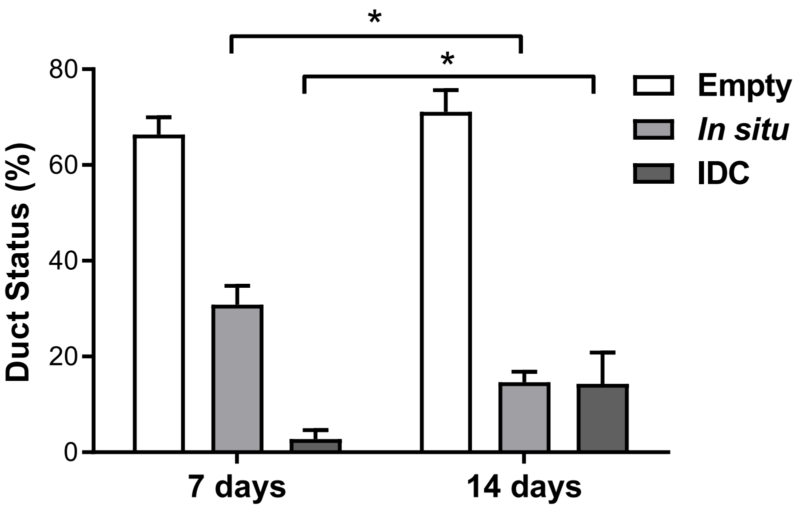

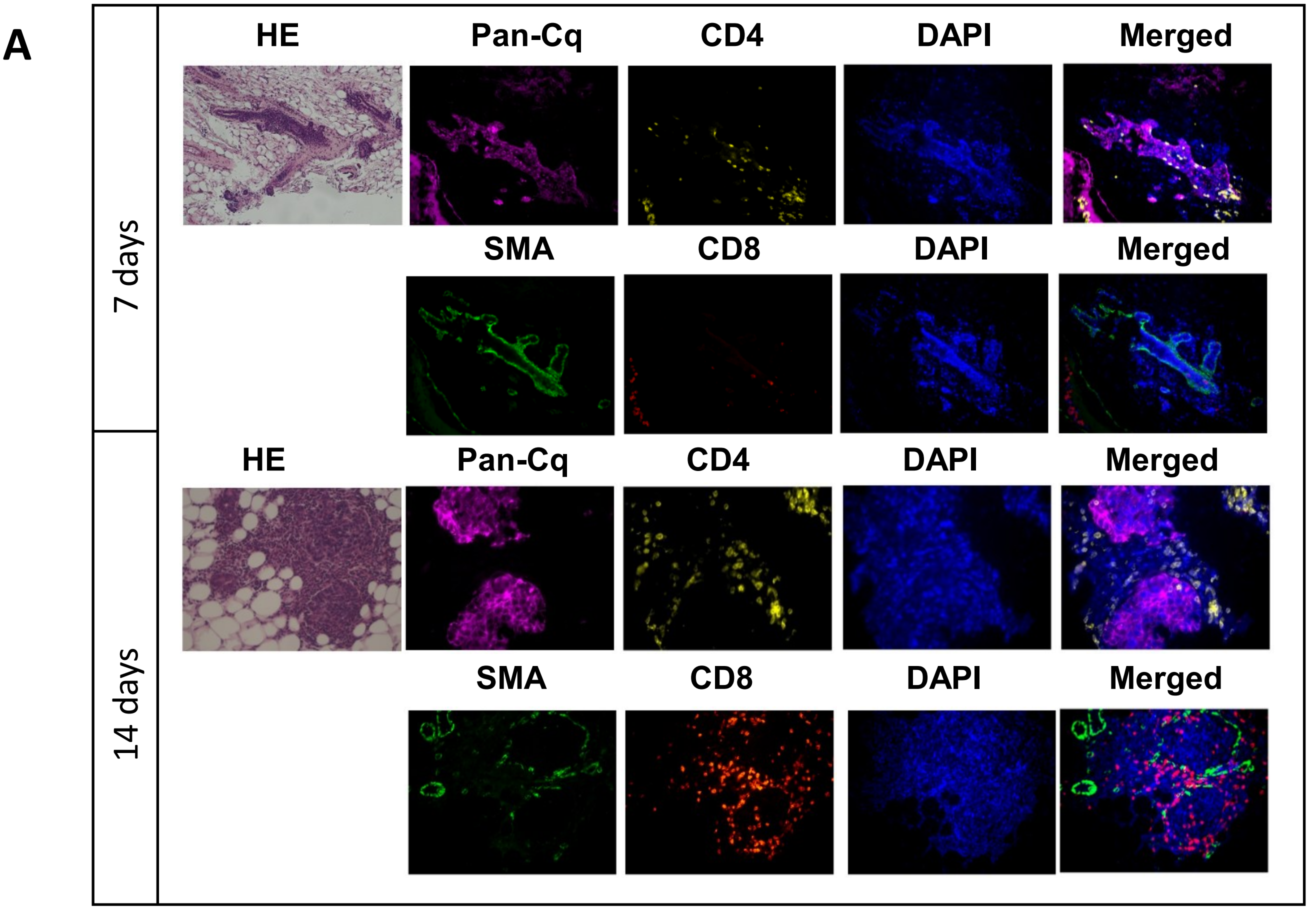

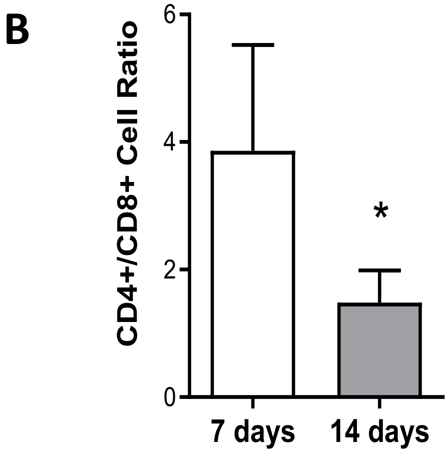

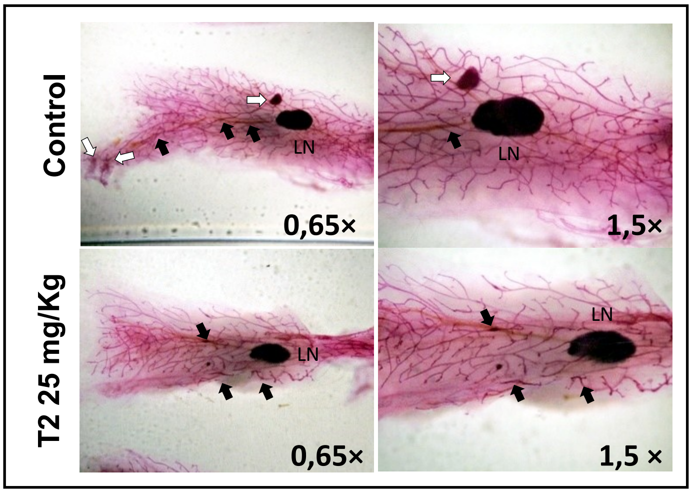
